# INTERLEUKIN-18 AS A THERAPEUTIC TARGET FOR WESTERN DIET-INDUCED CARDIOMYOPATHY

**DOI:** 10.1101/2025.09.23.678120

**Authors:** Pratyush Narayan, Salvatore Carbone, Adolfo G. Mauro, Eleonora Mezzaroma, Antonio Abbate, Stefano Toldo

**Affiliations:** Pauley Heart Center, Virginia Commonwealth University, Richmond, VA; Department of Physiology and Biophysics, School of Medicine, Virginia Commonwealth University, Richmond, VA; Nutrition Program, EVMS School of Health Professions, Division of Endocrine and Metabolic Disorders, Strelitz Diabetes Center, Old Dominion University, Norfolk, VA; Department of Pediatrics, Division of Pediatric Cardiology, Virginia Commonwealth University Medical Center, Richmond, VA; Robert M. Berne Cardiovascular Research Center, School of Medicine, University of Virginia, Charlottesville, VA, USA

**Keywords:** Interleukin-18, Systolic and Diastolic Function, Hyperglycemia, Obesity, Western Diet, Cardio-Immunology

## Abstract

**Background:** A diet high in saturated fats and sugars (Western diet-WD) promotes obesity and left ventricular dysfunction in the mouse, which is, at least in part, mediated by pro-inflammatory cytokine Interleukin-18 (IL-18). Therefore, we hypothesized that a blocking recombinant-murine IL-18 binding protein (IL-18BP) would rescue cardiac function in WD-fed mice.

**Methods:** In this 9-week study, 10-week-old adult C57BL/6J mice were assigned to standard diet (SD) or a WD. After 7 weeks of WD feeding, the mice were assigned to two groups: WD+IL-18BP (0.5 mg/kg daily, intraperitoneal injections) or WD control for the last 2 weeks of the study. Food intake, body weight, and glucose tolerance were assessed. Cardiac systolic and diastolic function were measured by Doppler echocardiography at baseline, 5 weeks, and 9 weeks. IL-18 plasma levels were quantified with ELISA.

**Results:** WD induced a significant increase in body weight, significantly worsened glucose tolerance, and significantly increased (worsening) in diastolic function (isovolumetric relaxation time -IRT- and myocardial performance index -MPI-) compared to SD. Rescue with IL-18BP in WD-fed mice resulted in a significant improvement in IRT and MPI, without significant changes in food intake, weight gain, or glucose tolerance.

**Conclusions:** IL-18BP rescued cardiac function in mice with WD-induced diastolic dysfunction, independent of weight gain and glucose tolerance. These results confirm the central and independent role of IL-18 in cardiac dysfunction associated with diet-induced obesity.

## INTRODUCTION

Heart Failure (HF) is clinically characterized by impaired cardiac function with symptoms of shortness of breath, dyspnea, fatigue, and poor exercise tolerance affecting 38 million people worldwide and 5.7 million people in the United States (Abbate et al. 2015; Braunwald 2013; Carbone et al. 2015; Carbone et al. 2017; O’Brien et al. 2014; Roh et al. 2022). Factors such as a diet rich in saturated fats and sugars (Western Diet -WD-), obesity, along with related metabolic derangements and inflammation, are widely implicated in HF (Abbate et al. 2020; Carbone et al. 2015; Carbone et al. 2017; Maurya et al. 2023; Van Tassell et al. 2012). A WD promotes both obesity and inflammation (Carbone et al. 2015; Carbone et al. 2017).

We have shown that the WD feeding in mice induces obesity and excess weight gain associated with progressive diastolic dysfunction over a period of 8 weeks. This diastolic dysfunction was partially reversible when the WD was interrupted and replaced by a standard chow diet (Carbone et al. 2015; Carbone et al. 2017). Additionally, we have shown that genetic deletion of Interleukin-18 (IL-18 KO mice) protected mice from cardiac dysfunction associated with WD-induced obesity, independent of body weight and glucose tolerance (Carbone et al. 2017; Carbone et al. 2018). In fact, IL-18 KO mice are more obese compared to their wild-type counterparts (Carbone et al. 2017; Netea et al. 2006), but did not develop systolic or diastolic dysfunction. However, whether the inhibition of this inflammatory pathway once the cardiac dysfunction develops is able to revert the diastolic dysfunction is unknown.

IL-18 is a pro-inflammatory cytokine that is part of the IL-1 family of cytokines, activated by the inflammasome pathway, including the NLRP3 inflammasome (NOD or NACHT, leucine-rich repeat (LRR), and pyrin domain (PYD)-containing protein 3) (Busch et al. 2021; Toldo and Abbate 2024). IL-18 binding protein (IL-18BP) is a naturally occurring IL-18 antagonist. The goal of this study was to target IL-18, with a recombinant murine IL-18BP (rmIL-18BP), to test its translational potential to rescue cardiac function in the setting of established WD-induced cardiac dysfunction. We hypothesized that in mice fed with WD, IL-18 drives cardiac dysfunction and that treatment with rmIL-18BP can improve it.

## METHODS

The experiments were conducted under the guidelines of laboratory animals for biomedical research published by the National Institutes of Health (No. 85-23, revised 2011). The study protocol (Fig. 1) was approved by the Virginia Commonwealth University Institutional Animal Care and Use Committee.

**Figure 1:**
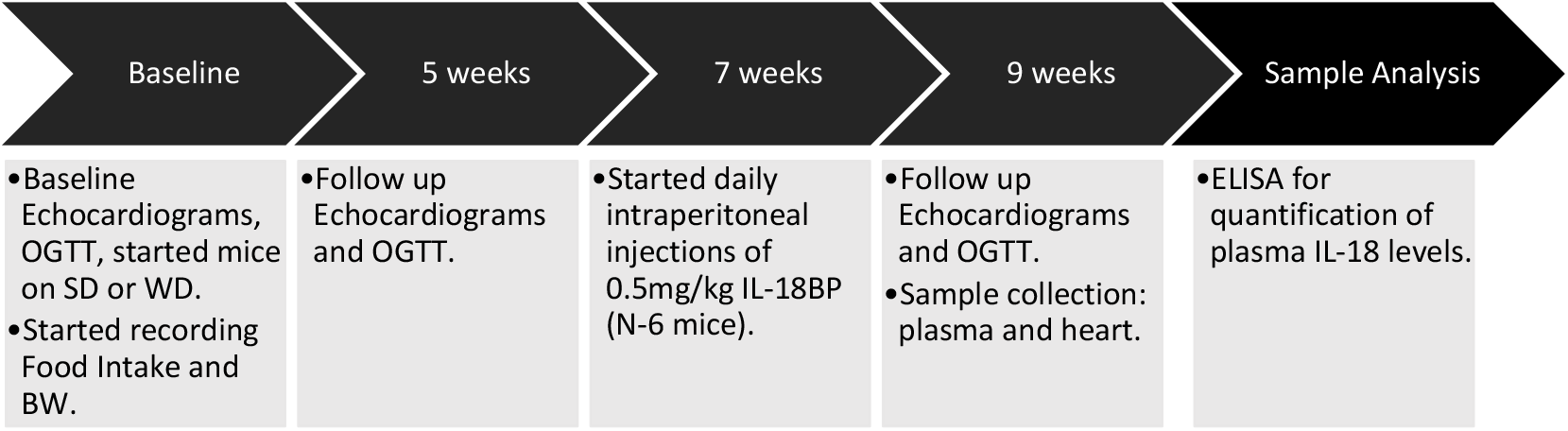
A visual timeline of our study design.

### Study design and procedures

We assigned 29 wild-type (WT) C57BL/J6 mice (Jackson Laboratories, Bar Harbor, ME), of adult age (10 weeks old) into three groups: SD group (fed standard diet for 9 weeks), WD group (fed Western Diet for 9 weeks), WD+IL-18BP group, which were fed WD for 9 weeks and started on IL-18BP daily injections starting at 7 weeks, for two weeks. All groups had an N≥6. The following diets were used: Standard chow diet (Teklad LM-485; Harlan, Madison, WI) and Western Diet (TD.88137; Harlan, Madison, WI). A comparison of the composition of these two diets is presented in Table 1.

**Table 1:** Diet composition. Comparison of the composition of the Standard diet, Western diet (Carbone et al. 2015; Cordain et al. 2005; Duffey and Popkin 2011; National Center for Health Statistics. 2016).

| Components in %kcal | Standard Diet | “Western Diet” |
| --- | --- | --- |
| Total fat | 17 | 42 |
| Saturated fat | 0.8 | 12.8 |
| Proteins | 25 | 15.2 |
| Total carbohydrates | 58 | 42.7 |
| Sucrose | 0 | 34 |
| Cholesterol | 0 | 0.2 |
| Sodium | 0.3 | 0.1 |
| Energy density (kcal/g) | 3.1 | 4.5 |

### Food intake and body weight

Food Intake (FI) was measured daily for the first two weeks of the study and later bi-weekly for the rest of the study. We expressed it as the average weekly food intake in each group of mice. Body weight (BW) was measured weekly.

### Doppler Echocardiography

All mice underwent pulse-wave Doppler (PWD) transthoracic echocardiography (TTE) using the Vevo770 imaging system (VisualSonic Inc, Toronto, Ontario and 30-MHz probe), at baseline (0 weeks), 5 weeks and 9 weeks, under light anesthesia with sodium pentobarbital at a dose of 30-50 mg/kg as previously described (Abbate et al. 2008; Respress and Wehrens 2010).

Briefly, the heart was visualized in a parasternal short-axis view. A 2-dimensional (2D) mid-ventricular position of the left ventricle (LV) was obtained at the level of the papillary muscles. Systolic parameters were measured in M-mode to quantify the end-diastolic and systolic parameters. The Teichholz equation was used to compute the ejection fraction (LVEF). PWD was used to measure isovolumetric contraction time (ICT), ejection time (ET) and isovolumetric relaxation time (IRT). Myocardial Performance Index (MPI), also known as the Tei index, was calculated for the LV using this equation: (ICT+IRT)/ET.

### Oral glucose tolerance test

An oral glucose tolerance test (OGTT) was performed at baseline, 5 weeks, and 9 weeks, with a 2-to 3-day interval between assessments, to allow for complete recovery from anesthesia. A glucose (Sigma Aldrich, St Louis, MO) solution was prepared at 20% volume per weight in water. It was administered at 2g/kg per mouse administered through an oral gavage needle (Becton Dickinson, Franklin Lakes, NJ). Glucose levels were measured prior to glucose administration (0 minutes), then at 15, 30, 60, and 120 minutes following the glucose load, using glucose strips and a portable glucometer (Clarity Diagnostics, Deerfield, FL). The area under the curve (AUC) was calculated from the glucose values.

### Treatment

The WD+IL-18BP rescue group received daily intraperitoneally (IP) injections of 0.5 mg/kg of lyophilized recombinant mouse IL-18BP (IL-18BP), His-tagged (IL-18BP-767M; Creative Biomart, Shirley, NY). The IL-18BP was diluted in normal saline prior to injections and administered from the week 7 to week 9 time points, using insulin syringes to minimize residual volume (0.5 ml; Becton Dickinson, Franklin Lakes, NJ).

### Sample collection

All mice were euthanized under deep anesthesia (sodium pentobarbital 70-100 mg/kg) followed by exsanguination at the 9-week time point. Whole blood was collected from the inferior vena cava (IVC) using 1cc tuberculin syringe and a 25-gauge 5/8-inch needle (Becton Dickinson, Franklin Lakes, NJ). Blood was collected in Vacutainer® PST Gel Lithium Heparin tubes (light-green top tubes; Becton Dickinson, Franklin Lakes, NJ). These tubes were stored on ice until being centrifuged at 2000 revolutions per minute (rpm) for 10 minutes at 4°C to separate plasma from the cells. The plasma was then transferred to 1.7ml Eppendorf tubes and snap frozen in liquid nitrogen, then stored at -80°C.

### IL-18 levels determination

Systemic levels of IL-18 were measured in the plasma using an enzyme-linked immunosorbent assay (ELISA) with a commercially available murine IL-18 ELISA kit sandwich assay (MBL International, Ina, Nagano, Japan). The optical density (OD) of the wells was measured at 450 nm with a microplate reader.

### Statistical Analysis

Statistical analysis and presentation of results were carried out with The R Project for Statistical Computing software (The Comprehensive R Archive Network) and RStudio IDE (Postit PBC, Boston, MA, USA) for Mac. The results have been presented as mean ± standard error of the mean. Variance between two or more groups was assessed by Levene’s test. Between-group differences with two groups were calculated with Student’s t-test (pooled variances) for unpaired data (equal variance). Between-group differences with three groups were calculated with one-way ANOVA followed by post-hoc Tukey HSD (equal variance) or Games-Howell (unequal variance) for pairwise comparisons between groups. Correlation was assessed via Pearson correlation and the strength and direction of the linear relationship between two variables was expressed as Pearson correlation coefficient (R). A *p-value* ≤ 0.05 was considered significant.

## RESULTS

### Effects of WD on IL-18 Plasma levels

After 9 weeks of study, the WD group had significantly increased IL-18 plasma levels compared to the SD group (158 ± 23 vs. 83 ± 19 pg/mL; P = 0.02) (Fig. 2A).

**Figure 2:**
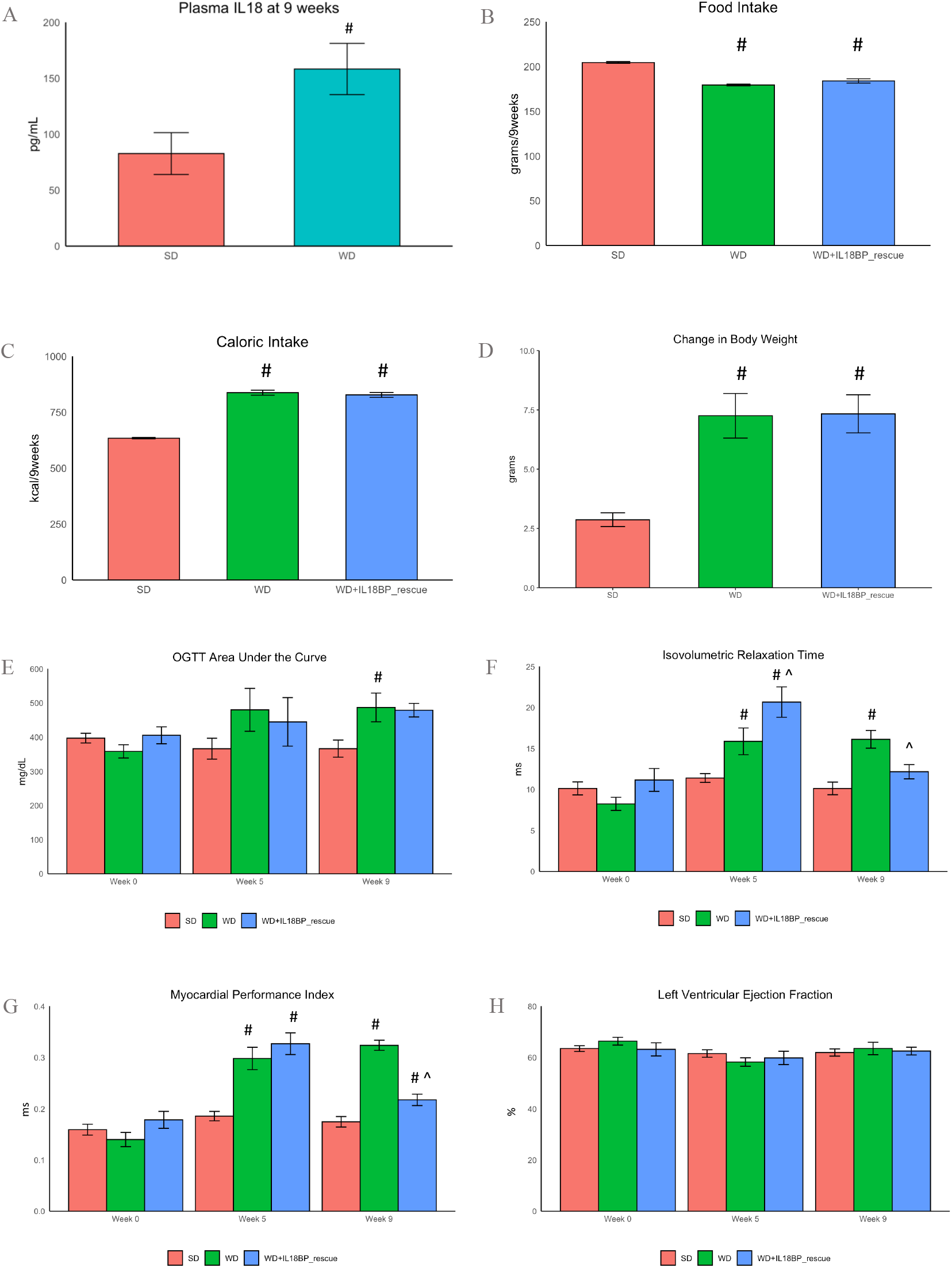
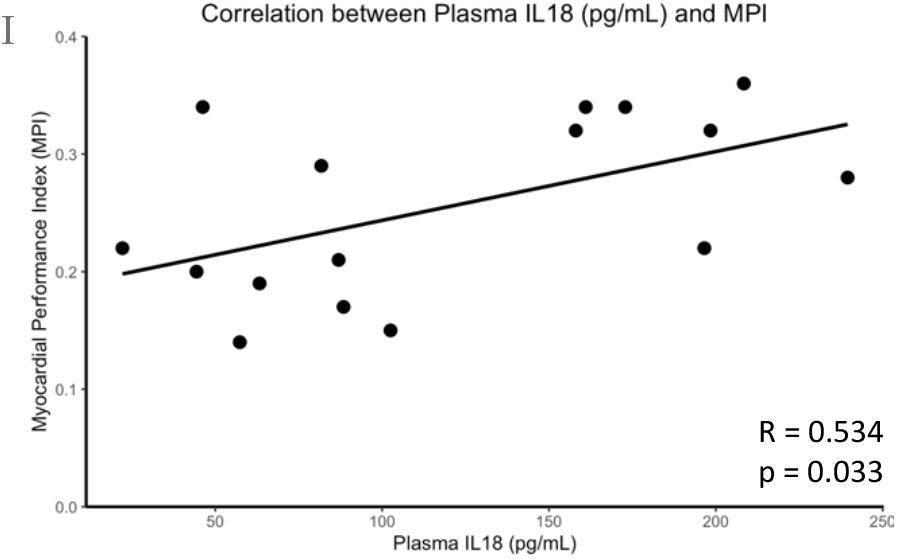
Western diet-induced inflammation, metabolic abnormalities, and cardiac dysfunction in mice fed with SD or WD. IL-18 plasma levels in the SD and WD groups, after 9 weeks of experimental time (A). Food intake in the SD, WD group, and the WD+IL-18BP rescue group after 9 weeks of feeding (B). Caloric intake in the SD, WD group, and the WD+IL-18BP rescue group after 9 weeks (C). Change in body weight in the SD, WD, and WD+IL-18BP rescue group, compared to the baseline body weight (D). Glucose tolerance was measured as the Area Under the Curve for the oral glucose tolerance test in the SD, WD, and WD+IL18BP rescue groups, at baseline, 5 (before IL-18BP treatment), and 9 weeks of feeding (E). Isovolumetric relaxation time (IRT) of SD, WD, and WD+IL18BP rescue group at baseline, 5 (before IL-18BP treatment), and 9 weeks (F). Myocardial performance index (MPI) was measured in SD, WD, and WD+IL18BP rescue groups at baseline, 5 (before IL-18BP treatment), and 9 weeks. (G). Left Ventricular Ejection Fraction (LVEF) measured at baseline, 5 (before IL-18BP treatment), and 9 weeks, in mice treated with SD, WD, and WD+IL-18BP rescue group (H). Correlation between plasma IL-18 levels and MPI in SD and WD group (I). # P ≤0.05 vs. SD; ^ P ≤0.05 vs. WD.

### Metabolic Effects of WD and Treatment with IL-18BP

At the end of the study, the weekly food intake (FI) (Fig. 2B) was significantly higher in the SD group (204±1.06 g) compared to the WD group (179±0.97 g; P ≤ 0.05) and WD+IL-18BP rescue group (184 ± 2.40 g; P ≤ 0.05). There was no significant difference in FI between the WD group and WD+IL-18BP rescue group (P = 0.1). However, the average mouse weekly caloric intake was significantly higher in the WD group (837 ± 11.22 kcal; P ≤ 0.01) and in the WD+IL-18BP rescue group (827 ± 10.78 kcal; P ≤ 0.01) compared to the SD group (634 ± 3.28 kcal) (Fig. 2C), as the WD has a higher energy density than the SD. There was no significant difference in caloric intake between the WD group and WD+IL-18BP rescue group (P = 0.8).

The weight gain (Fig. 2D) was significantly higher in the WD group (7.25 ± 0.94 g; P ≤ 0.01) and in the WD+IL-18BP rescue group (7.33 ± 0.80 g; P ≤ 0.01) compared to the SD group (2.87 ± 0.29 g). There was no significant difference in weight gain between the WD group and WD+IL-18BP rescue group (P = 0.9).

After 9 weeks, the AUC of the OGTT (Fig. 2E) was significantly higher in the WD group (487 ± 42 mg/dL; P = 0.03) compared to the SD group (366 ± 27 mg/dL). The AUC in the WD+IL-18BP rescue group was higher than the SD group but did not reach significance (479 ± 24 mg/dL; P = 0.1). There was no significant difference in AUC between the WD group and WD+IL-18BP rescue group (P = 0.9).

### Effects of WD and Treatment with IL-18BP on Cardiac Function

Cardiac diastolic function (Fig. 2F) was significantly impaired in the WD group, with higher IRT compared to the SD group, at 5 weeks (15.9 ± 1.6 vs. 11.4 ± 0.5 ms; P ≤ 0.02) and at 9 weeks (16.1 ± 1.1 vs. 10.1 ± 0.8 ms; P ≤ 0.01). Global LV function (Fig. 2G) was significantly impaired in the WD group, with higher MPI compared to the SD group at 5 weeks (0.30 ± 0.02 vs. 0.19 ± 0.01 ms; P ≤ 0.01) and at 9 weeks (0.32 ± 0.01 vs. 0.17 ± 0.01 ms; P ≤ 0.01).

Before being treated with IL-18BP, the WD+IL-18BP rescue group also had significantly higher IRT compared to the SD group (20.7 ± 1.9 vs. 11.4 ± 0.5 ms; P ≤ 0.01) and significantly higher MPI compared to the SD group (0.33 ± 0.02 vs. 0.19 ± 0.01 ms; P ≤ 0.01). However, treatment with IL-18BP initiated at 7 weeks promoted recovery in diastolic function in the WD+IL-18BP rescue group. At 9 weeks, the WD+IL-18BP rescue group had a significantly lower IRT compared to the WD group (12.2 ± 0.9 vs. 16.1 ± 1.1 ms; P ≤ 0.05) and significantly lower MPI compared to the WD group (0.22 ± 0.01 vs. 0.32 ± 0.01 ms; P ≤ 0.05). The improvement in IRT in the WD+IL-18BP rescue group was, to an extent, no longer significantly higher than the SD group at week 9 (12.1 ± 0.9 vs. 10.1 ± 0.8 ms; P = 0.3).

We did not detect any impairment in the systolic function with no significant difference in LVEF, between the groups of mice (Fig. 2H).

### Correlation between IL-18 Plasma levels and Cardiac Function

We found a significant positive correlation between the plasma levels of IL-18 measured at 9 weeks and the MPI (R=0.5; P=0.03) on the Pearson correlation test (Fig. 2I).

## DISCUSSION

In this study, we investigated a translational approach of treating mice with WD-induced cardiac dysfunction with pharmacological IL-18 blockade using recombinant mouse IL-18BP. We modelled this diet-induced obesity mouse using a WD with nutritional values similar to obesogenic diets often consumed in Western countries. We know that this type of diet is associated with elevated inflammatory markers and cardiac dysfunction, in humans and animal models, and specifically with diastolic dysfunction leading to heart failure with preserved ejection fraction (HFpEF) (Carbone et al. 2017; Cordain et al. 2005; Duffey and Popkin 2011). We also know that obesity and WD feeding in mice lead to an increase in IL-18,(Carbone et al. 2017). A chronic state of inflammation is an important but still not fully understood underlying factor in the pathophysiology of HF. For the first time in literature, we describe that WD-induced cardiac dysfunction is associated with elevated IL-18 levels and that targeting IL-18 with IL-18BP restores cardiac function after the establishment of WD-induced cardiac dysfunction.

IL-18, part of the IL-1 family of cytokines, is a cardio-depressant proinflammatory cytokine that induces cardiac dysfunction, both diastolic and systolic, and when it is blocked, reduces this dysfunction (Carbone et al. 2015; Dinarello et al. 2013). IL-18 has been associated with both structural and functional diastolic impairments (Boini et al. 2014; Carbone et al. 2017; Ehses et al. 2010). IL-18 levels are elevated in patients suffering from obesity and diabetes, both conditions that are associated with HFpEF (Carbone et al. 2017). Increased IL-18 in the serum of HF patients was associated with increased mortality (Abbate et al. 2015; Boini et al. 2014; Carbone et al. 2015; Carbone et al. 2017; Dinarello 2006; Netea et al. 2006; O’Brien et al. 2014). Additionally, comorbidities such as obesity and diabetes are often associated with diastolic dysfunction (Goldberg et al. 2005; Massie et al. 2008; Toldo and Abbate 2015; Wang et al. 2015).

The WD increases the mRNA levels of IL-18 in the heart and increases plasma levels of IL-18 (Carbone et al. 2017). Free fatty acids (FFAs) and simple carbohydrates (sugars), abundant elements of the WD, increase systemic levels of Interleukin 18 (IL-18) (Carbone et al. 2017). FFAs in the WD activate the IL-18 pathway and act as chronic stressors leading to low and chronic inflammation (Carbone et al. 2015; Regan et al. 2015). Toll Like Receptor 2 (TLR2) and TLR4, which are involved in the innate immune response, are activated by FFAs, contributing to local and systemic inflammation (Carbone et al. 2015; Ehses et al. 2010). These TLRs can lead to increased expression of pro-IL-18, which can be cleaved to its active form by the NLRP3 inflammasome (Carbone et al. 2017; Larsen et al. 2008; Marchington and Pond 1990; Netea and Joosten 2016; O’Brien et al. 2014; Valente et al. 2012; Yoshida et al. 2014).

Blockade of IL-18 has been studied in humans and mice using either rIL-18BP or an antibody targeting IL-18 (O’Brien et al. 2014). Blocking IL-18 in experimental studies has been shown to preserve or improve contractile function, decrease infarct size in acute myocardial infarction (AMI) and LPS-induced myocardial dysfunction animal models (O’Brien et al. 2014). In a model of chronic hypoxia, IL-18BP was used to block endogenous IL-18 and prevent the occurrence of diastolic dysfunction (Larsen et al. 2008). Blockade of IL-18 in the context of HF remains to be studied in humans.

WD-fed IL-18 knock out (IL-18KO) mice do not develop cardiac dysfunction despite having higher food intake and higher increase in body weight than WT mice on WD (Carbone et al. 2017). The total body deletion of IL-18 results in hyperphagia, leading to increased weight gain and impaired glucose tolerance (Carbone et al. 2017; Netea and Joosten 2016). IL-18KO mice fed with WD have lower myocardial fibrosis, as well as preservation of both diastolic and systolic function in IL-18KO compared to WT mice (Carbone et al. 2017). Therefore, the cardio-protection of WD-fed IL-18KO mice suggests a sufficient and causative role of IL-18 in WD-induced cardiomyopathy (Carbone et al. 2015; Carbone et al. 2017; Carbone et al. 2018). Although the KO models help us to investigate the mechanism, the clinical relevance of this type of approach is minimal.

In this study, we decided to use a translational approach by letting the mice develop diastolic dysfunction and try to reverse it by blocking IL-18. The dose of 0.5mg/kg of IL-18BP used in the treatment group is in the effective dose range that has been shown to work in a mouse model of arthritis (Banda et al. 2003; Dinarello et al. 2013; Faggioni et al. 2001; Plater-Zyberk et al. 2001). Interestingly, in the arthritis model, doses of 1 mg/kg and above lost effectiveness compared to 0.5 mg/kg. We found that, after the establishment of WD-induced diastolic dysfunction, IL-18BP treatment for 2 weeks reduced the IRT and MPI, close to normal levels, as shown by comparison with the same parameters in SD-fed mice. An increase the IRT is an indicator of diastolic dysfunction (Carbone et al. 2017). Similarly, an increase in the myocardial performance index (MPI) indicates a worsening of cardiac function, a lower efficiency of the cardiac cycle. In fact MPI is an index of the overall systolic and diastolic function of the left ventricle and therefore increase in MPI is an indicator of global cardiac dysfunction (Asami et al. 2019). Of course, the MPI must be taken with context of diastolic function and systolic function. Given, that in this study we did not see systolic impairment from LVEF measurement, we can safely state that the impaired MPI values was driven by the diastolic impairment measured with IRT.

When we correlated the MPI values of the mice in the SD and WD groups together, they had a significant positive correlation with the levels of plasma IL-18. Importantly, this is in accordance with our hypothesis that the levels of IL-18 drive the cardiac dysfunction, and further justifies the blockade of IL-18 as a strategy to treat or prevent it.

Unlike the cardiac function, treatment with IL-18BP did not affect the metabolic parameters in WD-fed mice. In our study, both the WD group and WD+IL-18BP rescue group had a significant weight gain at 9 weeks from baseline, compared to the SD group, which was associated with higher weekly energy/caloric intake (kcal). Additionally, WD induced glucose intolerance that was not influenced by IL-18BP treatment. This observation is important because, despite the neutral effects of IL-18BP treatment on metabolic parameters, it suggests that IL-18 blockade using recombinant IL-18BP does not induce more weight gain as previously observed by gene deletion in mice. (Carbone et al. 2017; Carbone et al. 2018; Netea et al. 2006).

This study, as any preclinical study, has inherent limitations. Our results should be viewed as preliminary findings into the attenuation of diastolic dysfunction using IL-18BP as a pharmacological blocker of IL-18 in an animal model. Our assessment of heart dysfunction was limited to echocardiography parameters and lacked more comprehensive hemodynamic measurements to estimate signs of HF, since we cannot assess other symptoms of HF in mice, such as dyspnea and fatigue, which can be assessed in humans. However, in prior studies, we have found that 8 weeks of WD induce an increase in LV end diastolic pressure, a sign of HF, and IL-18 deletion prevented this increase. Although heart function was improved in our treatment group, we cannot assess whether HF was completely rescued as we could not quantify the other symptoms which characterize HF. Future studies are needed to answer this question, and longer treatment with the WD and IL-18BP, could help to better characterize treatment limitations.

## CONCLUSION

In conclusion, the WD-induces cardiac dysfunction in the mouse correlates with IL-18 driven systemic inflammation. Exogenous IL-18BP administration after LV dysfunction is established promotes a recovery of the heart function.

## ACKNOWLEDGEMENTS

Mr. Narayan is funded by the C. Kenneth and Dianne Wright MD-PhD Scholar Award through the C. Kenneth and Dianne Wright Center for Clinical and Translational Research at Virginia Commonwealth University.

Dr. Abbate is supported by a NIH National Institute on Aging award (R01AG076360)

Dr. Abbate and Dr. Toldo are supported by an NIH-NHLBI award (R01HL174999)

